# The SUbventral-Gland master Regulator (SUGR) of nematode virulence

**DOI:** 10.1101/2024.01.22.576598

**Authors:** Clement Pellegrin, Anika Damm, Alexis L. Sperling, Beth Molloy, Dio S. Shin, Jonathan Long, Paul Brett, Andrea Díaz-Tendero Bravo, Sarah Jane Lynch, Beatrice Senatori, Paulo Vieira, Joffrey Mejias, Anil Kumar, Rick E. Masonbrink, Tom R. Maier, Thomas J. Baum, Sebastian Eves-van den Akker

**Affiliations:** The Crop Science Centre, Department of Plant Sciences, University of Cambridge, Cambridge CB2 3EA, UK; Department of Biochemistry and Metabolism, John Innes Centre, Norwich NR4 7UH, UK; Mycology and Nematology Genetic Diversity and Biology Laboratory, United States Department of Agriculture - Agricultural Research Service, Beltsville, Maryland, 20705, USA; Department of Plant Pathology, Entomology and Microbiology, Iowa State University, 2213 Pammel Dr., Ames, IA 50011, USA; Genome Informatics Facility, Iowa State University, 448 Bessey Hall, Ames, IA 50011, USA

**Keywords:** Plant-parasitic nematodes, Effectors, Effector regulation, transcriptional regulation

## Abstract

All pathogens must tailor their gene expression to their environment. Therefore, targeting host:parasite biology that regulates these changes in gene expression could open up routes to pathogen control. Here, we show that in the plant-parasitic nematode *Heterodera schachtii,* host signals (termed effectostimulins) within plant roots activate the master regulator *sugr1*. SUGR1, then, directly binds effector promoters, and orchestrates their production. Effector production, in turn, facilitates host entry, releasing more effectostimulins. These data show that gene expression during the very earliest stages of parasitism is defined by a feed forward loop for host entry. Importantly, we demonstrate that blocking SUGR1 blocks parasitism, underlining the SUGR1 signalling cascade as a valuable target for crop protection. Given that nematodes also parasitise humans and other animals, the potential impact is broad: disrupting effector production could, in principle, be applied to any pathogen that secrets effectors.

**Graphical abstract:** 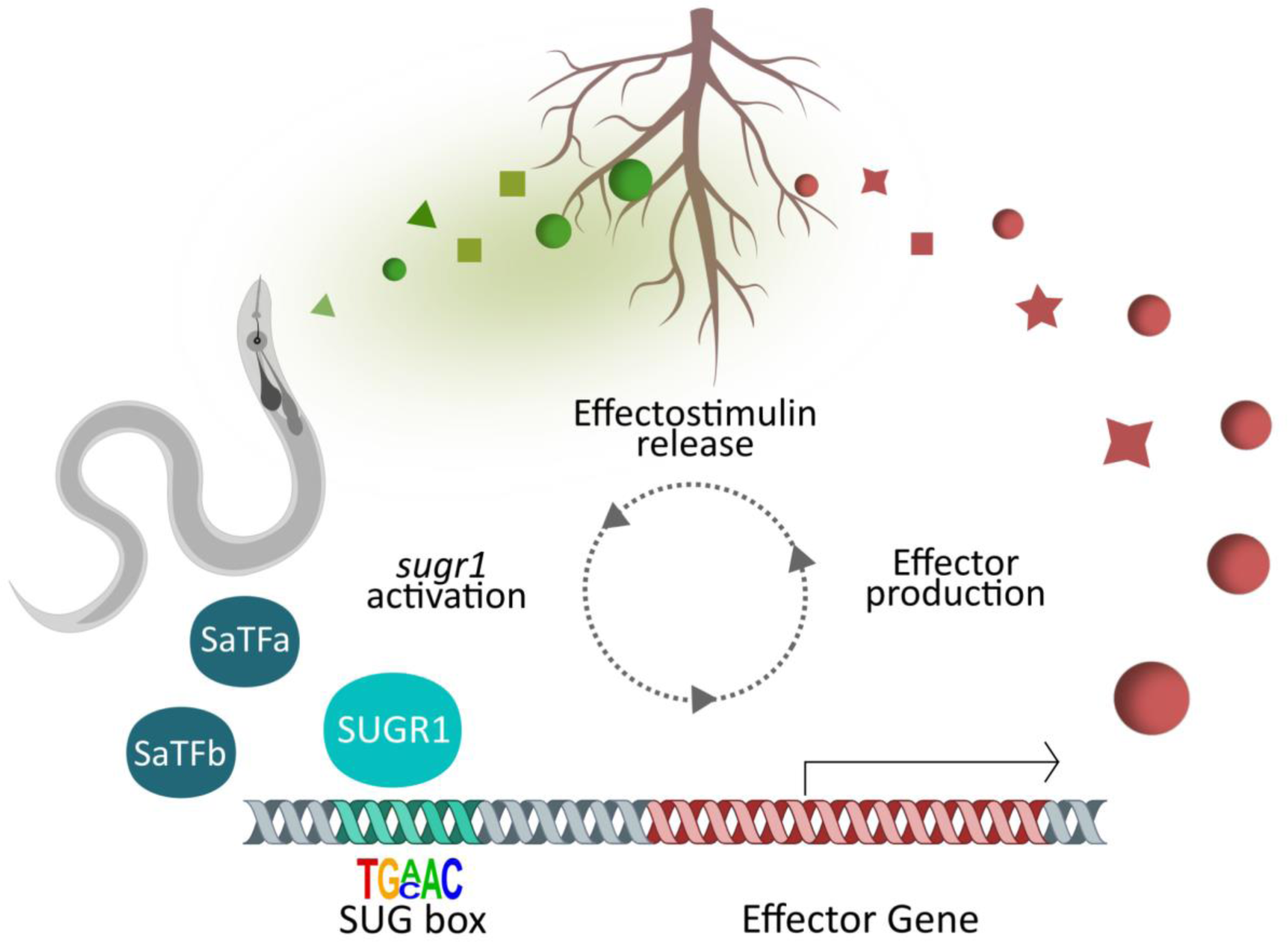

## Introduction

In humans, other animals, and plants, pathogen/parasite secretory/excretory products (often termed effectors) manipulate the host to benefit the invader (Win et al. 2012; Maizels, Smits, and McSorley 2018). These effectors can be recognised by the host, allowing the immune system to restrict infection, leading to an evolutionary arms race between host and pathogen (Wu, Derevnina, and Kamoun 2018; Perrigoue, Marshall, and Artis 2008). Effectors, and the host genes that they interact with, therefore, sit at the crux of engagement between kingdoms of life, defining disease or resistance.

Recent and rapid advances in effector biology have shaped our understanding of effector function and importance (Zhang et al. 2022). Effectors play pivotal roles during host invasion (Kubicek, Starr, and Glass 2014), immune suppression (Derevnina et al. 2021; Maizels, Smits, and McSorley 2018) as well as modulation of host physiology and development (Le Fevre et al. 2015) sometimes even culminating in the formation of novel organs (Siddique and Grundler 2018). As a result, various defence strategies aim to interfere with effectors, e.g. through recognition by resistance genes or by targeting effectors directly via RNAi (Prasad et al. 2019; Ali et al. 2017).

However, blocking the action of individual effectors will likely not lead to durable control of a given pathogen because effectors are at the interface of host-pathogen interactions and subject to intense evolutionary pressure. This has resulted in higher than background fixation of mutations compared to other genes (Eves-van den Akker et al. 2016), and the localisation of effectors in genomic regions associated with higher mutation rates (Sánchez-Vallet et al. 2018). This high rate of evolution, coupled with functional redundancy (Zheng et al. 2014) and overwhelming numbers (in some cases hundreds (Molloy et al. 2024)), impacts the robustness and practicality of effectors as targets for pathogen control. Indeed, resistance achieved through targeting effectors has been swiftly overcome (Brown 2015; Fouché, Plissonneau, and Croll 2018). A solution may be to target effectors indirectly by blocking the unifying aspects that regulate host:parasite biology during infection (Eves-van den Akker and Birch 2016).

Effectors production is precisely regulated in time and space to infect the host (Toruño, Stergiopoulos, and Coaker 2016; Nobori et al. 2020; Siddique et al. 2022; Yan et al. 2023). In this regard, all pathogens must recognise they are inside the host to effectively alter their physiology and gene expression. However, to the best of our knowledge, a signalling cascade from host cue to effector production has not been defined in a metazoan pathogen.

Here, we define such a signalling pathway in a cyst nematode - devastating pathogens of global agricultural importance that can cause yield losses of up to 90% in cereals (Nicol et al. 2011) and 80% in potatoes (Bernard, Egnin, and Bonsi 2017). We show that effector gene expression in the beet cyst nematode, *Heterodera schachtii,* responds to small molecule signals termed ‘effectostimulins’, which are found inside plant roots. When in contact with the nematode, effectostimulins activate the transcription factor SUGR1: a master regulator of effectors, including several known virulence determinants. SUGR1 is able to directly bind effector promoters and activate effector gene expression. We propose a model where, in a positive feedback loop, increased effector production facilitates host invasion, which in turn results in the release of more effectostimulins. Finally, we demonstrate that blocking this signalling cascade blocks parasitism, and translate these findings to the SUGR1 homologue in the soybean cyst nematode *Heterodera glycines*. This signalling cascade can be targeted on multiple levels (from blocking the host cues to blocking the regulator itself) and opens the door to analogous, novel control mechanisms in many pathosystems.

## Results

### Nematodes exposed to root extract are transcriptionally primed for infection

Most *Heterodera schachtii* effectors are maximally expressed after the nematode has reached the plant (Siddique et al. 2022). Therefore, we hypothesised that effectors, and indeed regulators thereof, might respond to plant signals. To simulate, and distinguish between, the perception of signals associated with host approach and host entry, we separated the molecules contained within roots (root extract), from those released into the rhizosphere (root diffusate (Figure 1A)) of the host *Sinapis alba*. Application of root extract and/or root diffusate altered the expression of 685 nematode genes, which were assigned to 6 distinct clusters (Figure 1B). Extract application, whether in combination with diffusate or alone, had the largest effect on gene expression (Figure S1), with 602 genes being assigned to the clusters “Extract up” or “Extract down” (Figure 1). Furthermore, the “Extract up” cluster was enriched in effectors (as predicted in Molloy et al. (2024)), secreted proteins, and gland cell expressed genes and contained the most effectors (48 effectors in “Extract up” vs 4 effectors in “All up” and 12 effectors in “Extract down” (Figure 1B)). Notably, all effectors in the “Extract down” cluster are expressed during later life stages.

**Figure 1:**
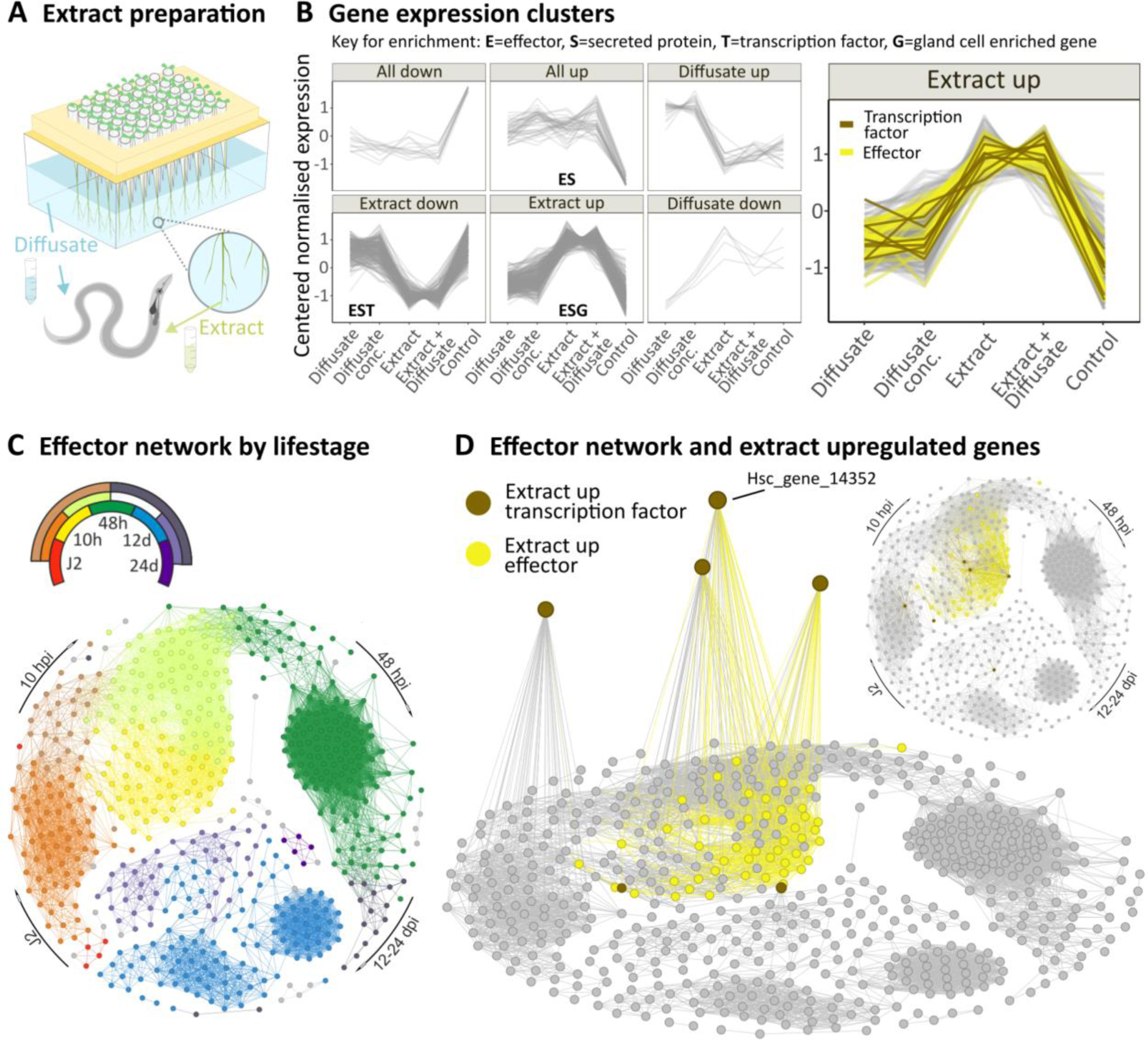
*Heterodera schachtii* gene expression responds to host cues. **A)** *Sinapis alba* plants were grown in tip boxes filled with water. *H. schachtii* second stage juveniles (J2s) were exposed to root diffusate (water) and/or extract prepared from the roots. **B)** Differential gene expression (|log2FC| ≥ 0.5 & padj ≤ 0.001) clusters that describe *H. schachtii* response to root diffusate and extract. Enrichment was determined in hypergeometric tests (p-value<0.05) **C)** Transcriptional effector network, where nodes represent effector genes predicted in Molloy et al. 2024 and edges represent correlations in gene expression across the nematode life-cycle of 0.975 or above (distance correlation coefficient). Colours indicate the nematode life stages. **D)** Transcriptional effector network highlighting effectors upregulated by *S. alba* root extract in yellow. Extract upregulated transcription factors with connections to the effector network (brown) are shown on the z axis where height is determined by connectedness to the effector network.

**Figure S1.**
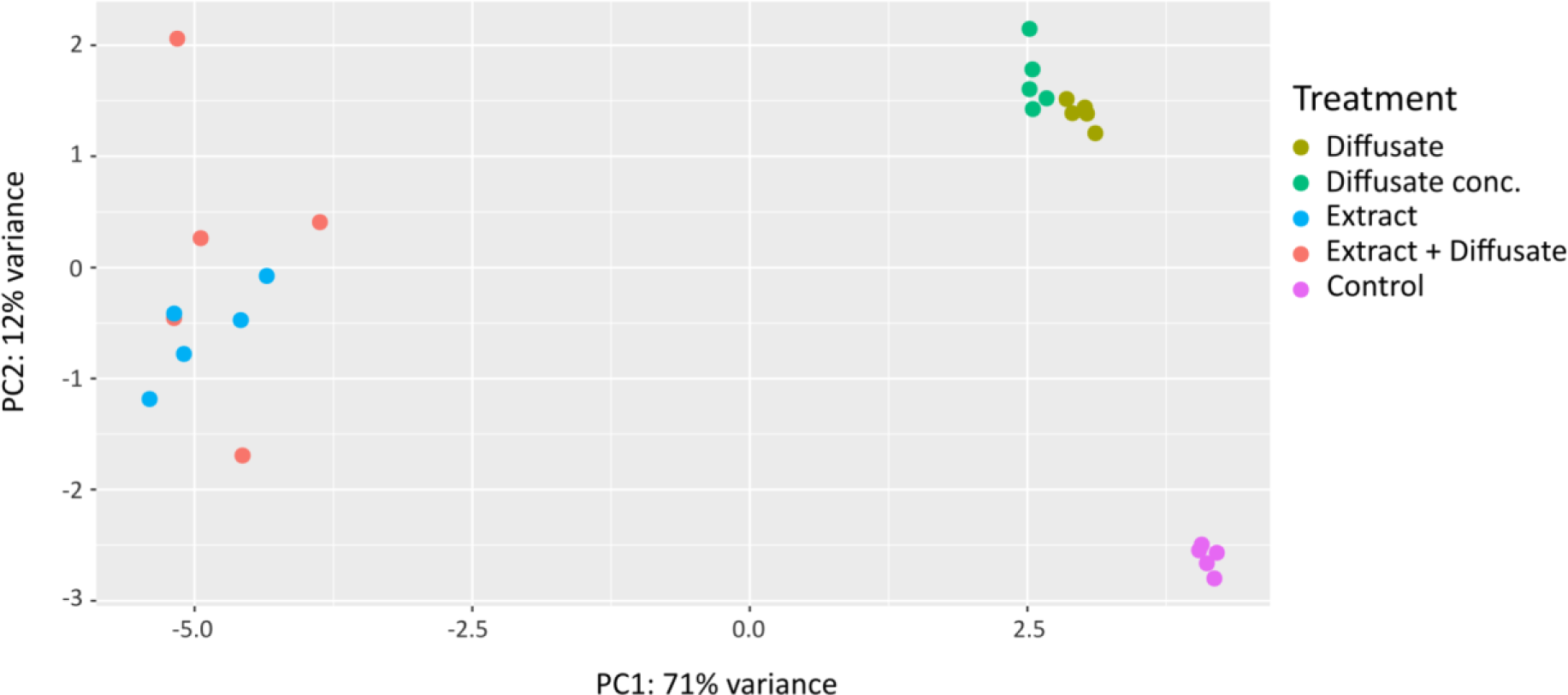
Principal Component (PC) Analysis of RNA sequencing data shown in Figure 1B.

Based on the hypothesis that the expression of a positive regulator of effectors is likely correlated with effector gene expression, we investigated the eight transcription factors also found within the “Extract up” cluster as putative regulators of effector gene expression. To prioritise these eight candidate regulators, they were cross referenced with the effector network (Figure 1C) described in Molloy et al. (2024). The effector network was constructed from independent life-stage specific transcriptome data from (Siddique et al. 2022) and displays the correlations of effector gene expression across the nematode life cycle. Of the eight candidate regulators, Hsc_gene_14352 is the most highly connected transcription factor to the network, and indeed the second most highly connected transcription factor of any kind. It is co-localised in the network with effectors expressed at the very earliest stages of host entry (measured 10 hours post infection (hpi)). Cross referencing the network with those effectors upregulated by root extract highlights this same time point (Figure 1D), independently validating the observation.

### Identifying the SUb-ventral Gland Regulator 1 (SUGR1)

Hsc_gene_14352 is a canonical nuclear hormone receptor, predicted to encode both a C-terminal DNA binding domain (DBD) and a predicted N-terminal ligand binding domain (LBD), and is expressed principally at 10 hours post infection (Figure 2A, B, and C). Nuclear hormone receptors are known to regulate a variety of processes (e.g. response to developmental, environmental and nutritional signals), and the family is expanded in nematodes (Taubert, Ward, and Yamamoto 2011). Nematode effectors are predominantly produced in two sets of gland cells. The two subventral gland cells are more active during the earlier stages of infection while the dorsal gland cell becomes active later in the nematode life cycle (Cotton et al. 2014). While Hsc_gene_14352 is reliably represented in targeted gland cell transcriptomic data of parasitic second stage juveniles (J2s (Molloy et al. 2024)), we used Sperling prep. fluorescence *in situ* hybridisation chain reaction (Sperling and Eves-van den Akker 2023) to show that Hsc_gene_14352 is specifically expressed in the subventral gland cells (Figure 1D). Taken together, these data show that Hsc_gene_14352 is expressed in the same cells, and at the same time, as subventral gland effectors.

Hsc_gene_14352 is, predominantly, a positive regulator of gene expression, as evidenced by comparative RNAseq analysis. Comparing gene expression in Hsc_gene_14352-silenced J2s against control *gfp*-silenced J2s reveals 297 differentially regulated genes (|log2FC| ≥ 0.5 & padj ≤ 0.001), the vast majority of which (77%) are concordantly down-regulated with Hsc_gene_14352 (Figure 3A and B). Consistent with functions of known subventral gland effectors (Molloy et al., 2024), the Hsc_gene_14352-regulon is enriched in GO terms (Table S1 and Figure S2) associated with carbohydrate metabolic processes (GO:0005975), polysaccharide catabolic processes (GO:0000272), and the parent term cellulose metabolic processes (GO:0030245). Indeed, Hsc_gene_14352 positively regulates 42 members of the predicted *H. schachtii* effectorome. Of those positively regulated effectors with experimental evidence of gland cell expression, 86% (18/21) are localised to the subventral gland, including several virulence determinants (Vanholme et al. 2007; Chen et al. 2005; Rehman et al. 2009; Long et al. 2013; Fanelli et al. 2014; Peng et al. 2016). Interestingly, among the activated genes encoding putative secreted proteins, 66 (61%) are not known members of the *H. schachtii* effectorome, highlighting potentially novel effectors (Figure S2). Therefore, to validate a subset of six activated genes, including known effectors and putative novel effectors, we confirmed subventral gland expression by *in situ* hybridisation (Figure 3C). Given the data, we have named the protein encoded by Hsc_gene_14352 the SUbventral Gland Regulator 1 (SUGR1).

**Figure 2:**
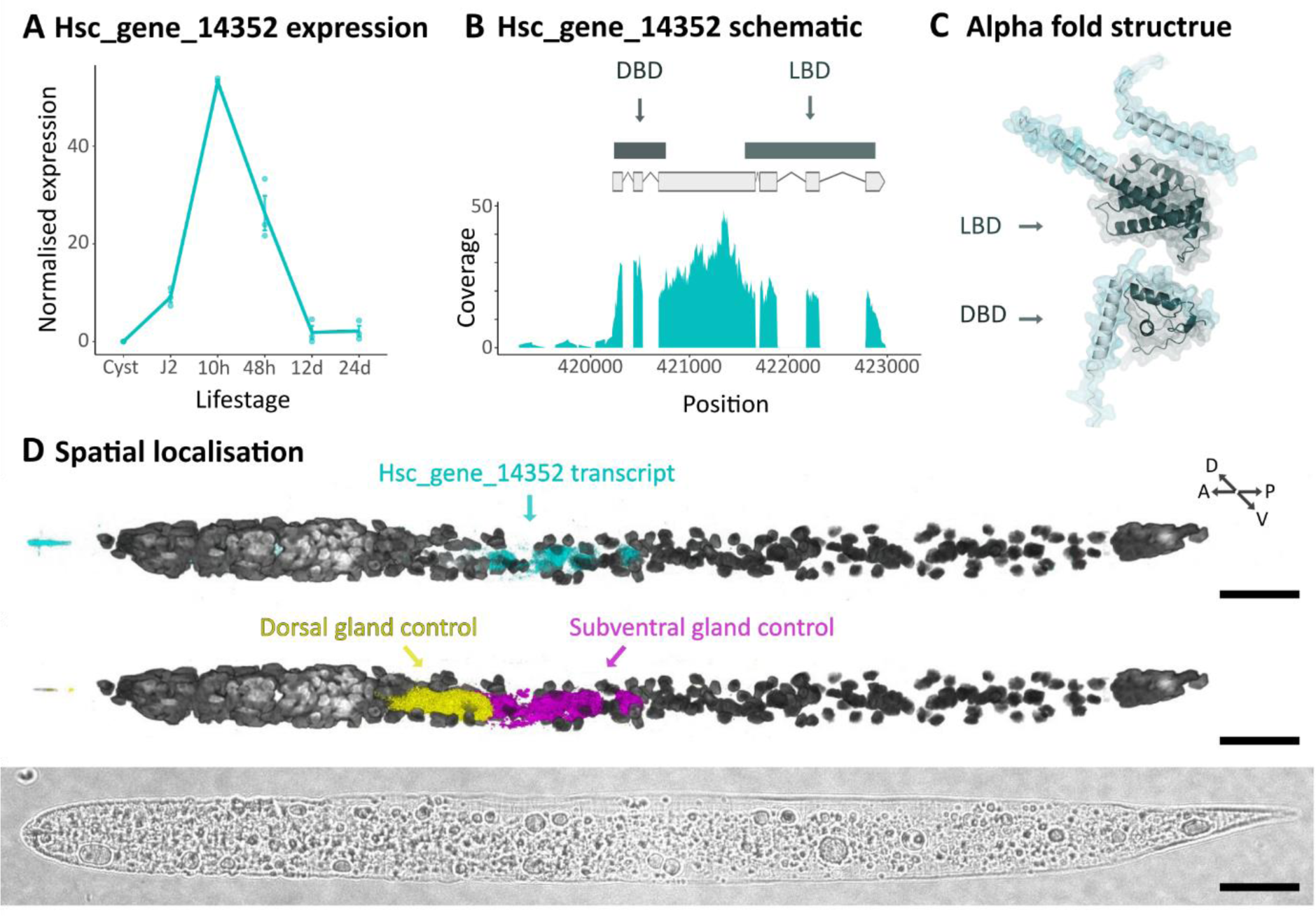
Characterisation of Hsc_gene_14352. **A)** Hsc_gene_14352 gene expression over the nematode life cycle. Data from (Siddique et al. 2022). **B)** Hsc_gene_14352 gene model, predicted to encode both a C-terminal DNA binding domain (DBD) and a predicted N-terminal ligand binding domain (LBD), and RNAseq coverage. **C)** Hsc_gene_14352 alpha fold structure **D)** Multiplexed Hybridisation Chain Reaction *in situ* for Hsc_gene_14352 transcripts (upper panel, cyan), compared to dorsal gland (Hsc_gene_2726) and subventral gland (Hsc_gene_21727 *eng2*) control transcripts (middle panel, yellow and magenta respectively). Nuclei stained with DAPI are shown in grey scale. Brightfield is shown in the bottom panel. Scale Bars represent 20 μm.

**Figure 3:**
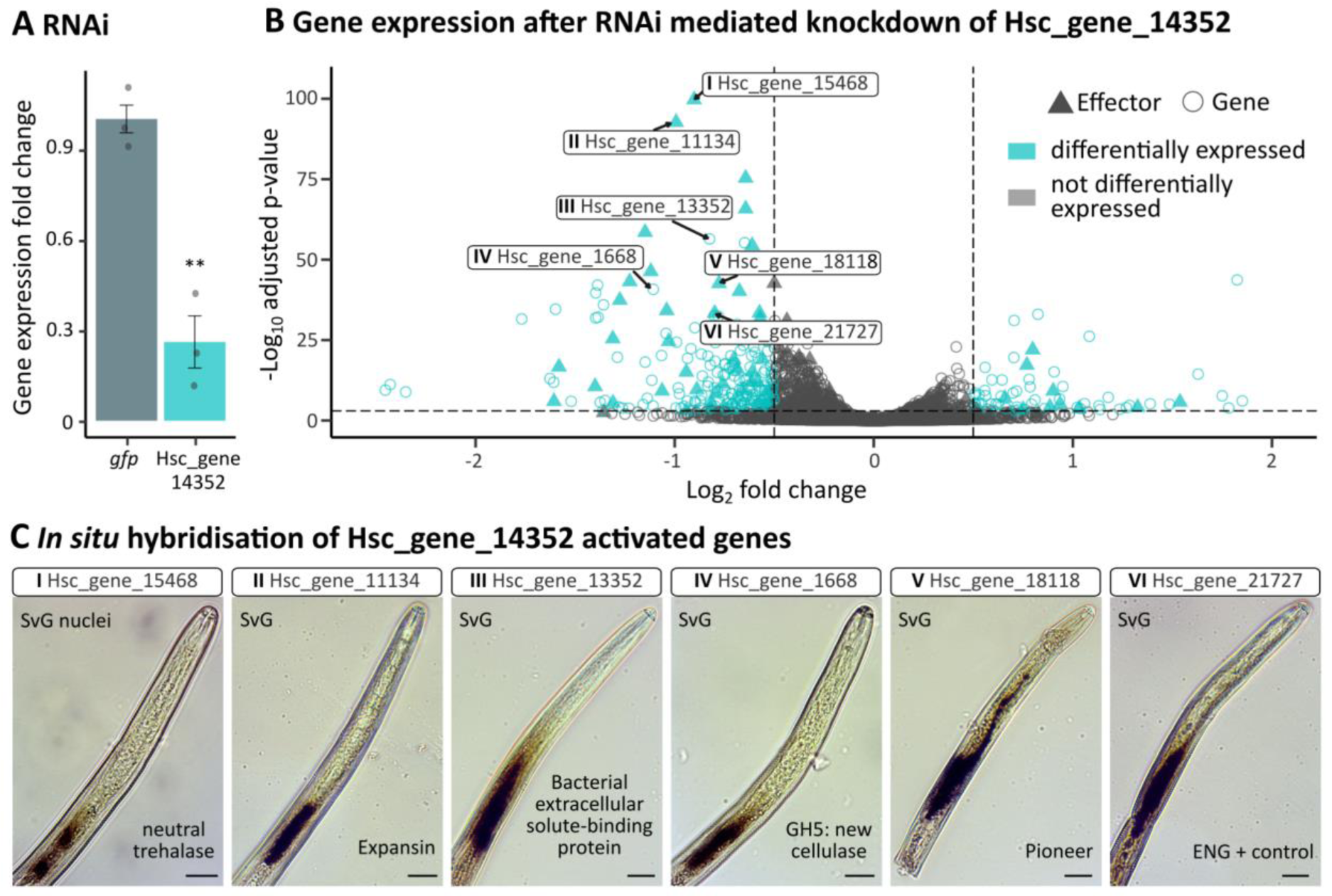
Hsc_gene_14352 is the SUbventral Gland Regulator 1 (SUGR1) **A)** Hsc_gene_14352 expression following RNAi mediated knockdown compared to a *gfp* control. Gene expression was determined by qPCR, normalised using the Pfaffl method and analysed using two sample t-test (p < 0.01=**). **B)** *Heterodera schachtii* gene expression following Hsc_gene_14352 knockdown vs. *gfp* control. Differentially expressed genes (|log2FC| ≥ 0.5 & padj ≤ 0.001) are highlighted in cyan. Effectors (as predicted in Molloy et al. (2024)) are triangles. Six (roman numerals) selected Hsc_gene_14352-regulated effectors/effector candidates are indicated. **C)** *In situ* hybridisation of six Hsc_gene_14352-regulated effectors/effector candidates. Scale bars represent 15 µm.

**Figure S2.**
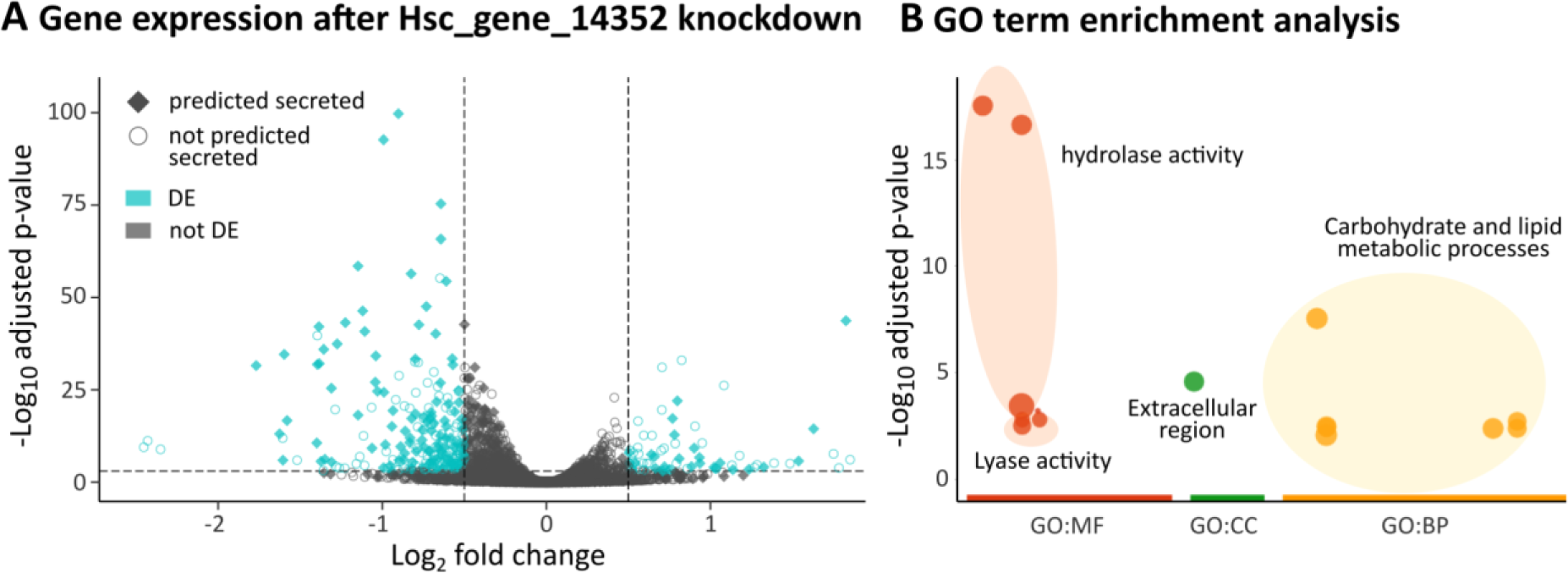
**A** *Heterodera schachtii* gene expression following Hsc_gene_14352 knockdown vs. *gfp* control. Differentially expressed genes (|log2FC| ≥ 0.5 & padj ≤ 0.001) are highlighted in cyan. Genes encoding proteins predicted to be secreted are diamond shapes. **B)** GO term enrichment analysis of Hsc_gene_14352 activated genes.

### Effectostimulins, small non-volatile signals inside plant roots, trigger a signalling cascade in Nematoda that upregulates effectors

*sugr1* is upregulated by root extract (Figure 1B), and SUGR1 in turn upregulates subventral gland effectors (Figure 3B). To characterise the earliest parts of this signalling cascade in more detail, we sought to determine whether root extract contains discrete activating signals. Removing molecules above 3 kDa from the extract, as well as heating to 45°C under a vacuum (as part of the extract preparation method), had no influence on the activating effect of the root extract (Figure 4A), implicating at least one small non-volatile signal. Removal of either strong anions or cations resulted in significantly reduced, but not abolished, activation of *sugr1* gene expression, perhaps indicating one or more charged signals (Figure 4B). Finally, and importantly, separating the contents of root extract based on their solubility in water, using High Performance Liquid Chromatography (HPLC), revealed multiple activating fractions in three activation peaks (Figure 4C). Taken together, these data can most easily be explained by a minimum of three discrete small molecule signals found inside plant roots that activate *sugr1* gene expression, and thereby effector expression: termed effectostimulins.

To characterise the later parts of the SUGR1 signalling cascade in more detail, we focused on the discovery that SUGR1 also controls the expression of three transcription factors: two SUGR-activated transcription factors (SaTF a and b) and one SUGR-repressed transcription factor (SrTF). Its ability to regulate effector gene expression may, therefore, be direct, indirect, or some combination thereof. We used Yeast-one-hybrid (Y1H) to determine whether SUGR1 and the SUGR-activated transcription factors can directly bind the 5’ proximal promoter regions of selected SUGR-activated effectors (up to 2kb upstream intergenic DNA), and therefore may have the capacity to directly regulate expression of these genes in cis. All three transcription factors are able to directly bind effector promoter regions in a partially overlapping manner, such that seven of the eleven tested effector promoter regions are bound by at least one transcription factor, and four are bound by all three in yeast (Figure 4D, Figures S3 and S4). Interestingly, SUGR1 also directly binds the promoter region of SaTFb in yeast (Figure 4D, Figures S3 and S4).

SUGR1-regulated interactions in cis are likely mediated by a conserved DNA motif (Figure 4E). Remarkably, in all of the following cases, using a differential motif discovery algorithm on the promoters of SUGR1-regulated effectors, subventral gland effectors, and early-stage effectors from the J2-10 hours post infection supercluster (as defined in (Siddique et al. 2022)) identifies a homologous sequence. Comparison between the enriched motifs reveals a conserved “core” of TG[C|A]AC, which is also the reverse complement of a canonical nuclear hormone receptor binding site (Weikum, Liu, and Ortlund 2018; Vivanco Ruiz et al. 1991). This motif is termed the SUventral Gland box (SUG box) following the established convention (Eves-van den Akker et al. 2016).

Together with our understanding of effectostimulins, these data paint a SUGR1-centric network of interactions that underlies the upregulation of effectors in the subventral gland cells at the very earliest times of host infection, based on host-derived signals.

**Figure 4:**
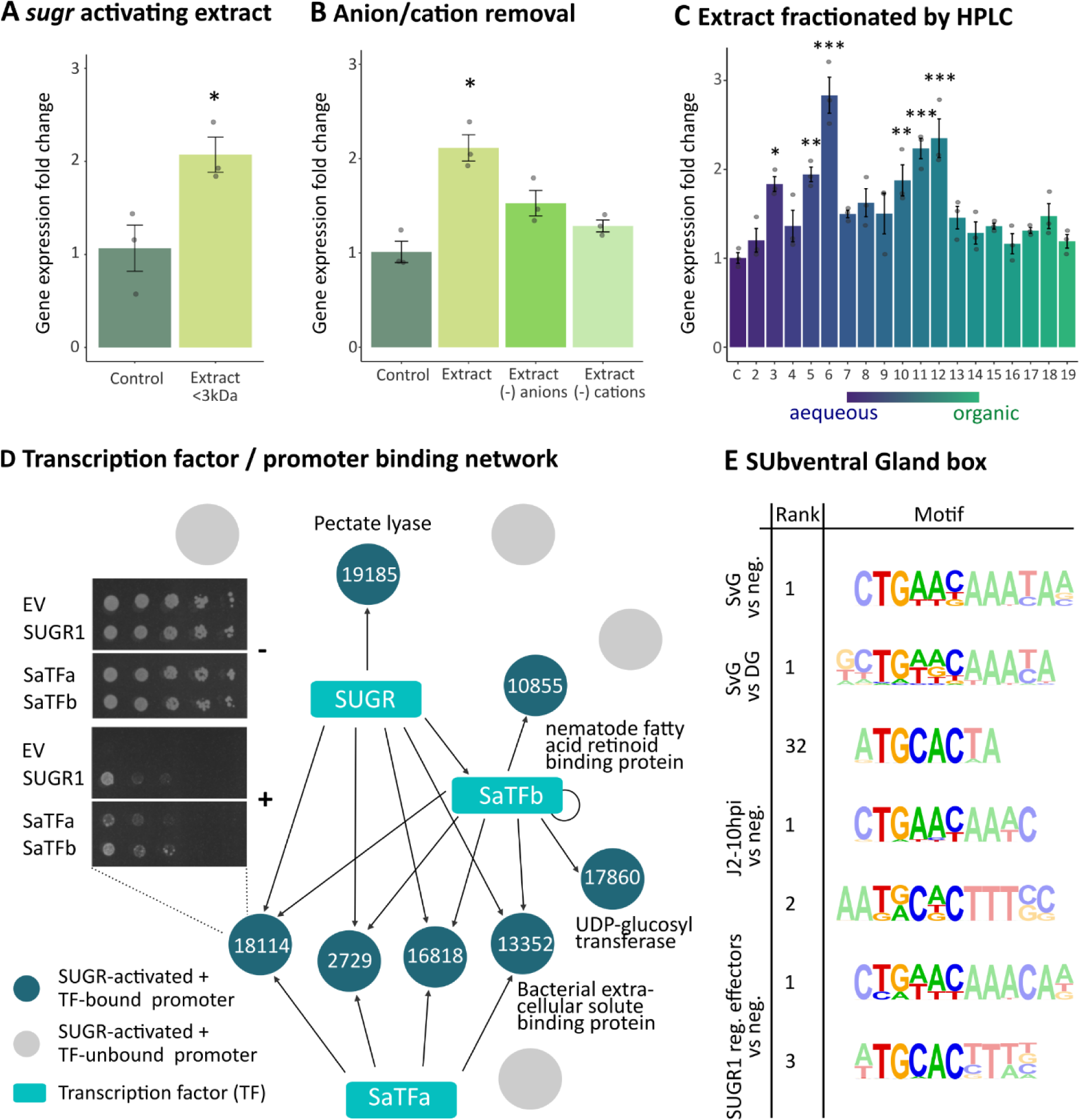
A signalling cascade regulating nematode effector production. **A)** Effect of *Sinapis alba* root extract on *sugr1* gene expression in *Heterodera schachtii* as determined by qPCR and analysed by two-sample t-test. **B)** Effect of ion depleted root extract on *sugr1*. Data were analysed using Kruskal-Wallis and Dunn’s test. **C)** Effect of *S. alba* root extract fractions (fractionated by High Performance Liquid Chromatography) on *sugr1*. Data were analysed using one-way ANOVA and Tukey HSD (Honestly Significant Difference) test. For A-C data were normalised using the Pfaffl method. Asterisks indicate treatments statistically significantly different to the water control (p<0.05=*; p<0.01=**; p<0.001=***) **D)** Network of transcription factor:promoter (up to 2 kb upstream intergenic DNA) interactions. Promoters bound by at least one transcription factor in yeast-one-hybrid screens (Figure S3-4) are highlighted in dark green. **E)** DNA motif associated with subventral gland effectors (SvG), SUGR1 regulated effectors, or effectors found in the J2-10 hpi supercluster from (Siddique et al. 2022). Motif enrichment analyses were performed with the respective promoters (800 bp from start codon) compared to a negative control set (neg.) or dorsal gland effectors (DG).

**Figure S3:**
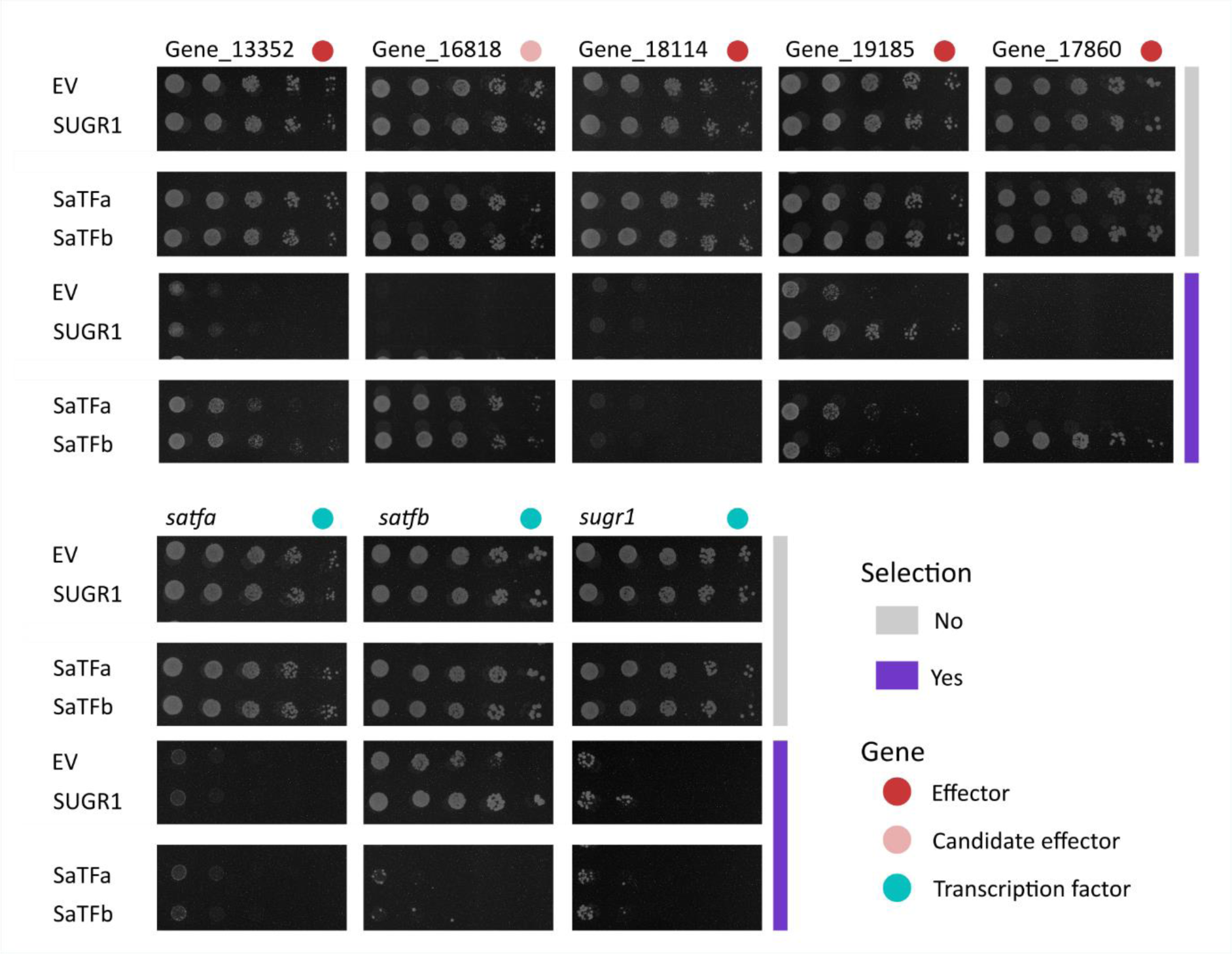
Yeast-one-hybrid pictures for full length promoters. Binding of SUGR1, SaTFa, and SaTFb to promoters (up to 2kb upstream intergenic DNA) of SUGR1-activated genes (Figure 3B). Promoters of effector genes (as predicted in Molloy et al. 2024 and/or validated via *in situ* hybridisation (Figure 3C) are highlighted with red circles. Yeast were grown with (purple) or without (grey) Aureobasidin A selection and yeast growth compared to the empty vector control (EV). Pictures were cropped but all comparisons for the same promoter come from the same plate.

**Figure S4:**
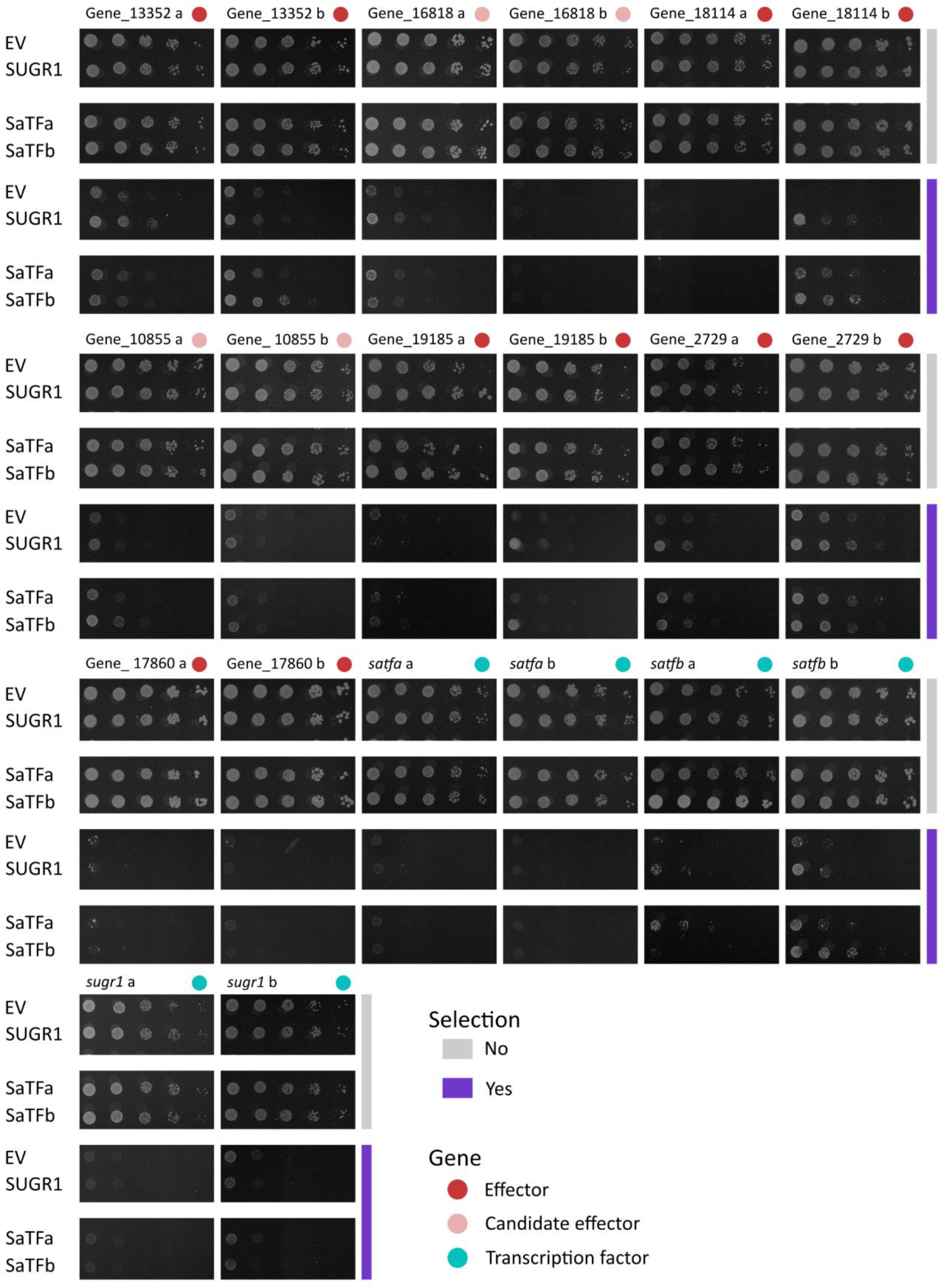
Yeast-one-hybrid pictures for promoter halves. Binding of SUGR1, SaTFa, and SaTFb to promoter halves of SUGR1-activated genes (Figure 3B) which were additionally analysed in two parts (proximal (b) and distal (a) halves). Promoters of effector genes (as predicted in Molloy et al. 2024 and/or validated via *in situ* hybridisation (Figure 3C) are highlighted with red circles. Yeast were grown with (purple) or without (grey) Aureobasidin A selection and yeast growth compared to the empty vector control (EV). Pictures were cropped but all comparisons for the same promoter come from the same plate.

### Blocking SUGR1 blocks parasitism

Given that SUGR1 positively regulates 42 effectors, several known virulence determinants, and multiple plant cell wall degrading enzymes, we hypothesised that blocking SUGR1 would impair the process of plant colonisation. Specifically, the very earliest stages of host entry are likely perturbed, given the dominant role of the subventral glands at this time. We therefore tested whether RNAi-mediated silencing of *sugr1* would also impact the ability of *H. schachtii* J2s to invade young mustard seedlings. Silencing *sugr1* significantly reduced the number of J2s observed inside the root compared to the *gfp*-silenced control (1.32 +/− 0.23 *sugr1* vs 6.7 +/− 0.78 *gfp*) after 10 hours of infection (Figure 5A). This confirms the involvement of SUGR1 in host colonisation and, given that a moderate reduction of *sugr1* expression is amplified to a much larger reduction in pathogenicity, highlights the importance of SUGR1 signalling in general. Taken together, these data suggest that the SUGR1 signalling cascade is a valuable target for crop protection: stopping the nematode infection before it enters its host.

*Heterodera schachtii* is both an economically important pathogen in its own right, and the model cyst nematode. Its close sister species, the soybean cyst nematode *H. glycines,* is the most economically important cyst nematode globally and the most damaging pathogen of any kind to US soy production (Savary et al. 2019). Importantly, these species have a common origin of parasitism, and remarkable conservation in effector repertoire (93% of *H. schachtii* effectors have a homolog in *H. glycines* (Molloy et al. 2024)). Given their relatedness, we were able to identify a single unambiguous homolog of *sugr1* in *H. glycines* (Hetgly00282, 80% identical across 100% query coverage, and only three amino acids different in the DNA binding domain). Knockdown of *H. glycines sugr* (*Hg sugr*) resulted in a concomitant knockdown of three out of four canonical subventral gland effectors tested (a pectate lyase, Hetgly20776; an expansin, Hetgly05367; and a gycosyl hydrolase 53 arabino-galactanase, Hetgly14426 - Figure 5B). Analogous to SUGR1, HgSUGR acts predominantly as a positive regulator, activating the expression of 57/86 regulated genes (Figure S5). Furthermore, proteins predicted to be secreted are significantly enriched (about 1.6 times more than expected) in HgSUGR activated genes (as determined in hypergeometric tests; p-value<0.05), and 58% of the GO terms associated with HgSUGR activated genes are also found for SUGR1 activated genes (Table S2). Importantly, knockdown of *Hg sugr* resulted in a significantly reduced number of *H. glycines* J2s per root (Figure 5C). Taken together, these data suggest that SUGR1 and HgSUGR regulate similar processes.

**Figure 5:**
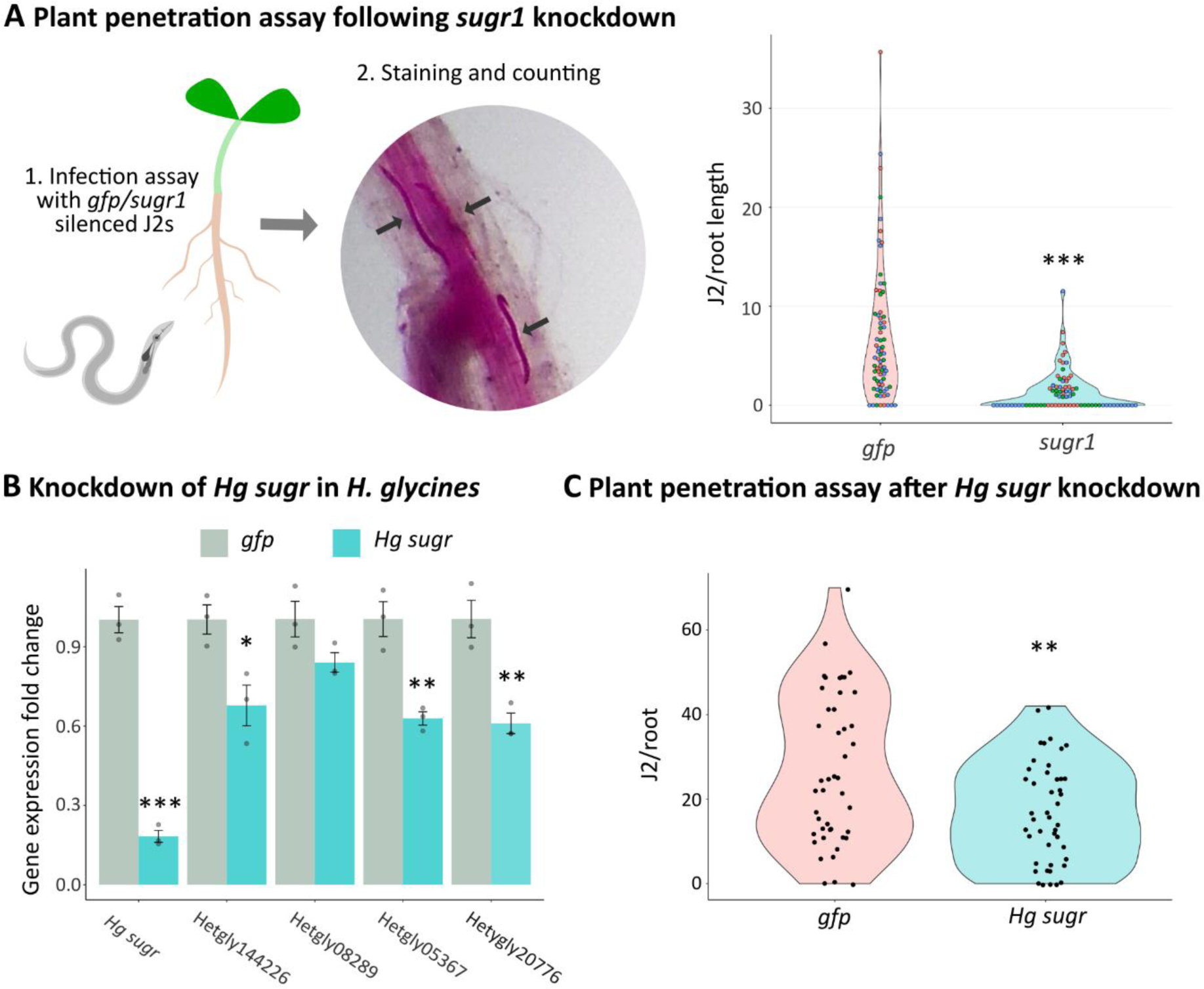
SUGR is required for full pathogenicity. **A)** *Sinapis alba* plants were infected with *Heterodera schachtii* second stage juveniles (J2s) following RNAi mediated silencing of *sugr1* or *gfp* (control). The impact on nematode parasitism was determined by J2s per root area. Asterisks indicate a significant difference compared to the *gfp* control at FWER<0.001 (Games-Howell test). **B)** *H. glycines sugr* (*Hg sugr*) was silenced in the same manner, and the effect on gene expression of it and four corresponding canonical subventral gland effectors was tested by qPCR and data normalised using the Pfaffl method. Asterisks indicate significantly different treatments compared to the respective *gfp* control (p<0.05=*; p<0.01=**; p<0.001=***; two sample t-test). **C)** Plant penetration assay with *H. glycines* J2s on *Glycine max*, following RNAi mediated silencing of *Hg sugr* or *gfp* (control). Asterisks indicate a significant difference compared to the *gfp* control at FWER<0.01 (Games-Howell test).

**Figure S5:**
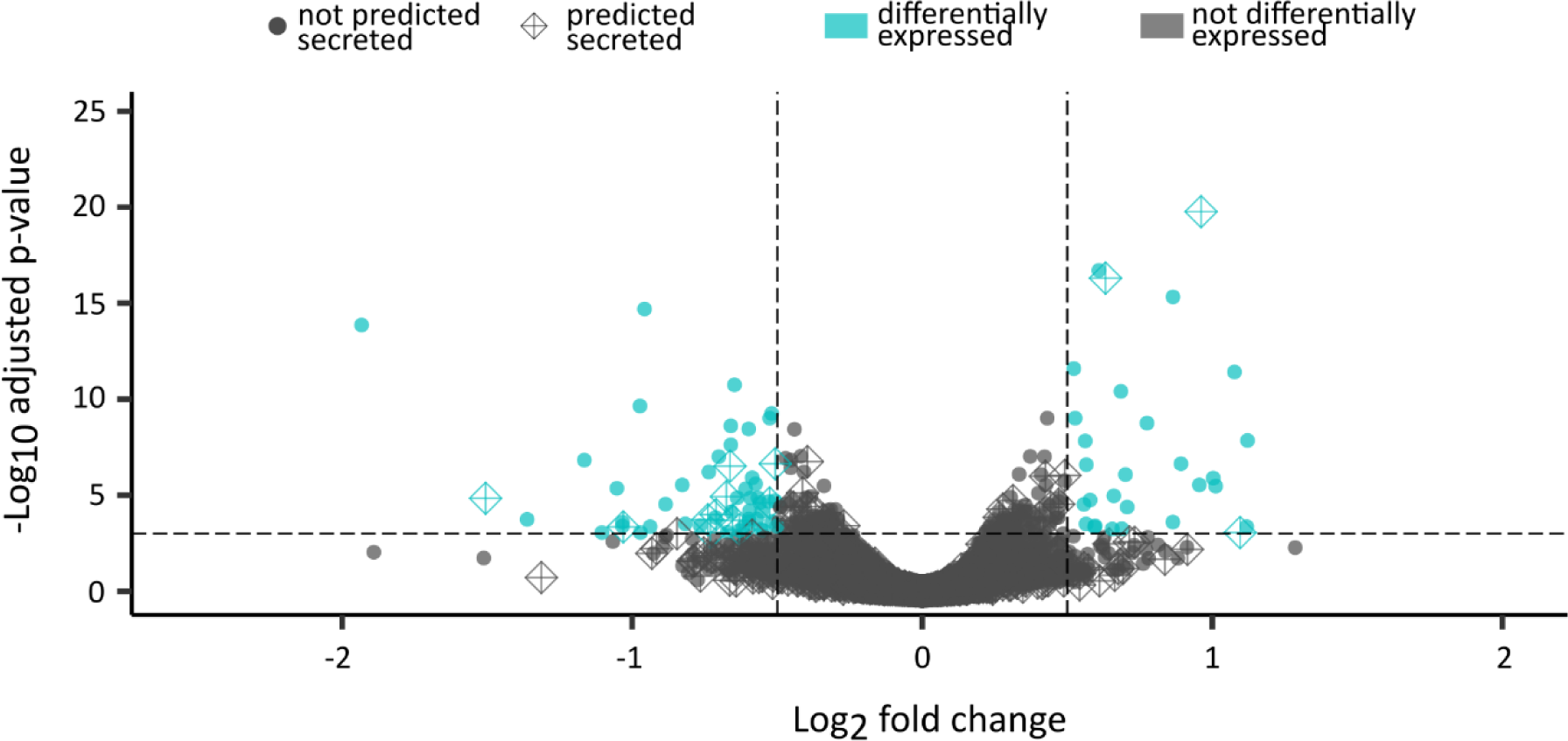
*Heterodera glycines* gene expression following *Hg sugr* knockdown vs. *gfp* control. Differentially expressed genes (|log2FC| ≥ 0.5 & padj ≤ 0.001) are highlighted in cyan. Genes encoding proteins predicted to be secreted are diamond shapes.

## Discussion

### Effectostimulins

All pathogens must tailor their gene expression to the environment in which they find themselves. In the case of cyst nematodes, this is strikingly evident in light of their biology. Juvenile worms have extremely limited energy reserves: they can remain dormant in the egg for decades, hatch, locate and enter the host, migrate through host cells, and establish a feeding site – all without feeding (Wyss and Zunke 1986). Effectostimulins - signals found *inside* plant roots that drive the expression of effectors - fulfil a fundamental requirement to spare resources when the nematode must, and promote parasitism when it counts.

We posit that effectostimulins must be distinct from host attraction signals because maximal effector expression before reaching the host (i.e. during the attraction phase) would be prohibitively wasteful. Effectostimulins encode the information that a permissive site within the host has been reached. Therefore, they must be present in, but not necessarily descriptive of, the host. They must be conserved enough to encode this information to the invader, so that it can be relied upon for such profound changes in pathogen physiology, gene expression, behaviour etc. Indeed, reliability of the signal may partly explain that effectostimulins of the *H.schachtii:S.alba* pathosystem are redundant, non-volatile, charged, small molecules. These data, together with our assertions above, will inform hypotheses on the nature of the signals and the future experiments to identify them.

Although the molecules are almost certainly distinct, effectostimulins are nevertheless a generalisable concept to other, if not all, pathosystems. Indeed, the authors are unaware of a pathosystem that does not alter its gene expression during infection. Recent studies in other plant-parasitic nematodes (Bell et al. 2019; Teillet et al. 2013; Duarte et al. 2015), and other eukaryotes (Wu et al. 2020), can be viewed through this lens, and further support generalisability.

### A feed forward loop for host entry

The data presented allow us to build a conceptual model for effector regulation in this system. Effectostimulins contained within plant cells are released upon the very earliest stages of host probing with the nematode stylet. They then activate the transcription factor and master regulator *sugr1*, which in turn (directly and likely indirectly), orchestrates the production of effectors (including many cell wall degrading enzymes). Effector production likely leads to increased cell penetration (Rehman et al. 2009), releasing yet more effectostimulins, which trigger even more effector production. This model is therefore, by its nature, a feedforward loop for host entry.

We showed that there are at least three discrete Effectostimulins and that SUGR1 and SUGR1-activated transcription factors bind effector promoters in a partially overlapping manner. The presence of multiple activating signals and transcription factors may perform two, not necessarily mutually exclusive, functions. Firstly, they may fine-tune the production of effectors depending on the host/environment in ways we do not yet fully understand. Alternatively, or in addition, such redundancy in the signalling cascade of both signals and transcription factors may increase robustness of the system to a variable host/environment. Nevertheless, in spite of this redundancy, we demonstrate that disrupting only one component, in this case SUGR1, is sufficient to disrupt the system.

**Figure 6.**
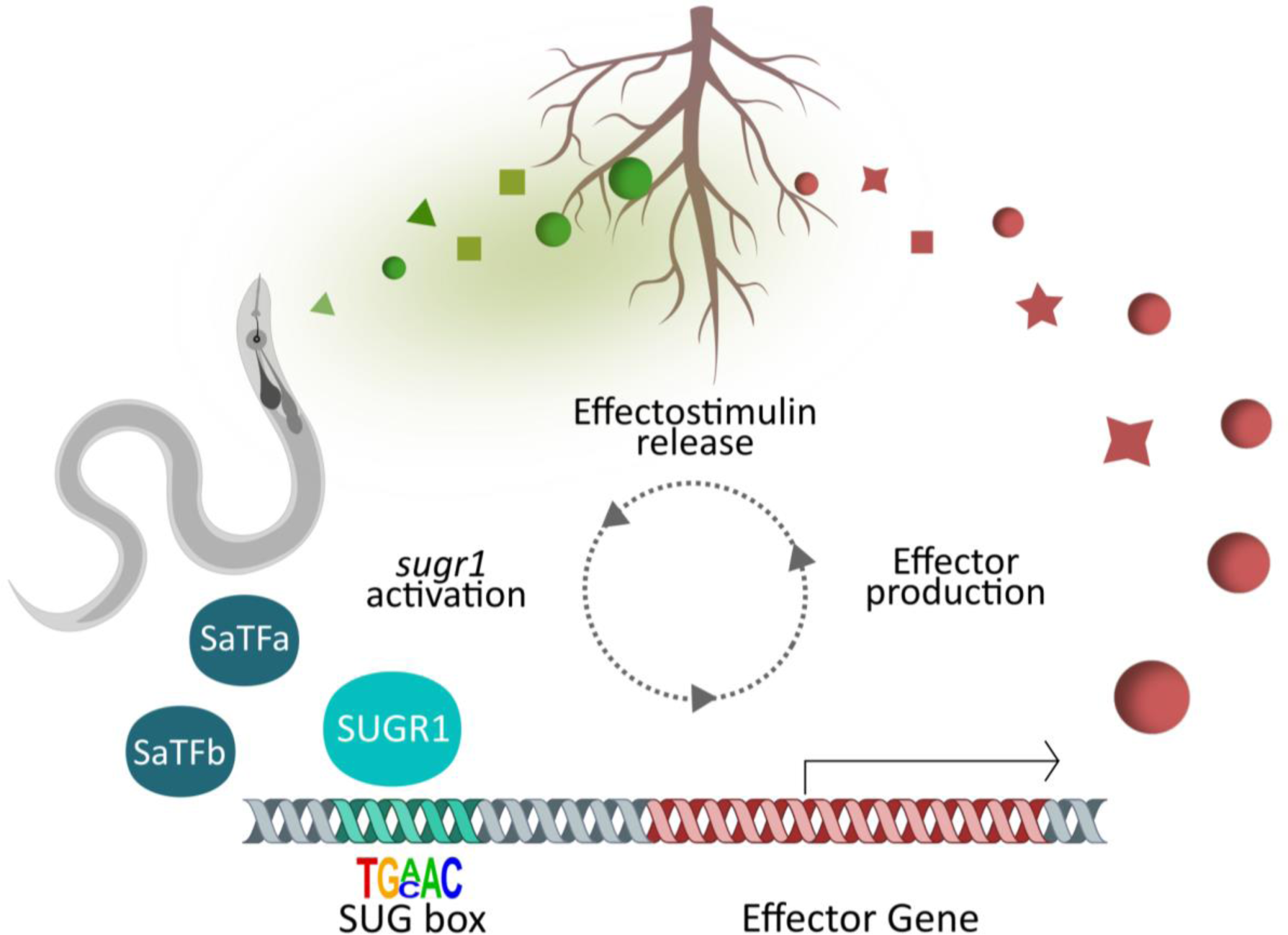
A feed forward loop for host entry. Effectostimulins in plant roots (green shapes) are released in the earliest stages of plant invasion. They then activate the master regulator *sugr1* (cyan), orchestrating the production of effectors, including many cell wall degrading enzymes (red shapes) likely via the SUG box. Effector production, in turn, leads to increased cell penetration, releasing more effectostimulins, triggering even more effector production.

### Future identification of other regulators and virulence determinants

Effectors sit at the crux of engagement between kingdoms of life. Here, we identify the first gene of any kind, shown to be required for effector production in any plant-parasitic nematode. Identification of further effectors and their regulators will help elucidate these central players of inter-kingdom communication. Among the SUGR1-activated genes encoding putative secreted proteins, 61% are not known effectors. Understanding the regulators of effectors may therefore represent a novel method of effector discovery in this system. In addition, four of the top eight most connected transcription factors to the effector network in Molloy et al. (2024) are also found in the “Extract up” cluster. These findings validate our approach and suggest nematode regulators of effector production in other glands and/or at other times of infection are within reach.

For other eukaryotic pathogens, we have some understanding of both positive and negative regulators of effector production. Positive regulators include orthologues of the transcription factor Wor1 that have been implicated in virulence of several phytopathogenic fungi, often regulating developmental aspects but also effector production (Michielse et al. 2009; Santhanam and Thomma 2013; D. W. Brown, Busman, and Proctor 2014; Mirzadi Gohari et al. 2014; Okmen et al. 2014; Tollot et al. 2016) and orthologues of Pf2, first shown to regulate effectors in the fungus *Alternaria brassicicola* (Cho et al. 2013; Rybak et al. 2017; Jones et al. 2019; See and Moffat 2023; Clairet et al. 2021; John et al. 2022). Negative regulators of effector gene expression prior to plant infection include Rgs1 of *Magnaporthe oryzae* (Tang et al. 2023). Despite these intriguing and valuable insights into the regulation of pathogen effector production, a missing piece of the puzzle is the integration into a signalling cascade starting with activating host cues. A holistic model, building on the identification of Effectostimulins, could extend the potential impact associated with an understanding of Effectostimulins yet further.

More progress in understanding effector production and utilisation for pathogen control has been made in the field of bacteriology. An emerging idea in bacteriology is therapeutics that block virulence factors, so called anti-virulence drugs. These novel strategies are proposed to reduce the selective pressure, since bacteria are not directly killed, and alleviate the problem of antibiotic resistance (Dehbanipour and Ghalavand 2022). Indeed, substances that target the Type III secretion system (Sharma, Elofsson, and Roy 2020) or effector secretion (Aburto-Rodríguez et al. 2021) have been identified.

### Routes to application

The discovery of SUGR1 as a *perturbable* master regulator of nematode virulence opens the door to promising novel routes to nematode control, targeting effector production instead of individual effectors. Blocking the nematode internal machinery that regulates effector production is promising because: i) blocking the production of effectors blocks all associated effectors at the same time; and ii) the machinery involved in effector production is hidden from the plant immune system and not genetically primed for evolution, likely leading to more durable resistance (Eves-van den Akker and Birch 2016). Future research will focus on testing the efficacy of targeting other components of the feed forward loop, and driving impact in crop protection.

The understanding of Effectostimulins themselves provides opportunity. For example, modulating Effectostimulin metabolism by conventional breeding, CRISPR/Cas genome editing (Jhu, Ellison, and Sinha 2023), or even root microbiome engineering (Korenblum et al. 2020), are all in principle plausible. Effectostimulins could also be applied in the field to undermine resource management and exhaust the nematodes prematurely. The finding that SUGR1 is a nuclear hormone receptor opens further doors for novel control mechanisms. Nuclear hormone receptors are often bound by ligands resulting in activation, repression, or relocation. Identifying the ligands of SUGR1, which may or not be synonymous with Effectostimulins, could therefore lead to further opportunities to disrupt the system. Nuclear hormone receptors are unusual in that they are considered “druggable” transcription factors (Weikum, Liu, and Ortlund 2018) due to their ligand binding domain, which makes screening for substances that block SUGR1 directly eminently feasible.

Cyst nematodes belong to the economically most damaging plant-parasitic nematode, they are the dominant nematode threat in UK/northwest Europe, and the number one pathogen in soybean (Savary et al. 2019). Control measures are extremely limited and the few nematicides available are being successively removed from the market (Price et al. 2021). The discovery of SUGR1 as a master regulator of nematode virulence opens the door to novel, much needed methods for nematode control, targeting the effector production instead of individual effectors. Importantly, we show disrupting SUGR signalling likely disrupts similar processes across the genus, potentially extending the applicability to other important agricultural pests. Regardless, the theory of blocking effector biogenesis is generalisable. Taken together, these findings promise to expand our toolkit in ensuring global food security. Moreover, nematodes are not only capable of agricultural and ecological catastrophes, but can also parasitise humans and other animals via the secretion of effectors. This context highlights immense potential impact: disrupting effector production could, in principle, be additionally applied to the fields of human and veterinary medicine, or indeed any pathogen that secrets effectors.

## Material and methods

### Common material

*Sinapis alba* (cv. albatross), and *Heterodera schachtii* populations “Bonn”, originally from Germany, (as per the reference genome (Siddique et al. 2022) and “IRS”, originally from The Netherlands, were used in this study. For the yeast-one-hybrid screen the *Saccharomyces cerevisiae* Y1HGold strain (Takarabio) was used. For bacterial transformation chemically competent cells of the *Escherichia coli* strain DH5α were used. All primers used are available in Table S3.

### Effectostimulin extraction and associated analyses

#### Extraction

*Sinapis alba* seeds were sterilised with 20 % bleach solution (Parazone) for 20 min and grown on wet filter paper at 21 °C for seven days. Alternatively, to separate Effectostimulins inside and outside roots, plants were grown in pipette tip boxes filled with 200 ml sterile, ultrapure water. To collect root diffusate, the water was exchanged after 7 days and collected 48 hours later. To prepare extract, roots were ground in ultrapure water (0.5g/1ml), centrifuged at 10,000 rpm for 2 min and the supernatant was collected. For size exclusion, the extract was centrifuged in vivaspin columns (<3kDa MWCO; Cytiva) at 4 °C. For ion removal, Pierce strong ion exchange columns (ThermoFisher) were used following supplier’s instructions. If needed, the extract was concentrated using a Concentrator plus (Eppendorf) at 45 °C until all liquid was removed and resuspended in the required volume. To generate data shown in Figure 1, Extract (<3kDa) was concentrated 3:1 to resemble biological conditions. Diffusate was used non concentrated (Diffusate) and concentrated 5:1 (Diffusate conc.). For the ion exchange (Figure 4B) and fractionation (Figure 4C) experiments the extract was concentrated 6:1 to achieve high activation levels. In the size exclusion experiment (Figure 4A), extract was prepared from 5-day old *S. alba* plants, the centrifugation step was replaced by filtration (0.45 µm), and the extract concentrated ten times.

#### Application to nematodes

*Heterodera schachtii* cysts were obtained from infected sand (Stichting IRS), isolated using sieves (4000, 2000, 500, 125, 63 microns) and transferred to hatching jars (Jane Maddern Cosmetic Containers). Hatching was induced by 3 mM Zinc chloride solution, jars kept at 21°C and J2s were collected every 2-3 days. At least 15,000 J2s per replicate were treated with 50 µl of *S. alba* root extract, *S. alba* root extract fractions or *S. alba* root diffusate at 21 °C and 700 rpm for 4 h. As a control, 50 µl of sterile, ultrapure water were added instead. Subsequently, nematodes were flash frozen in liquid nitrogen and stored at −80 °C.

#### RNA extraction and qPCR

Frozen *H. schachtii* J2s were ground to powder in a Geno/Grinder 2010 (Spex Sample Prep) in three 30 s long cycles at 1200 strokes/min. Subsequently, total RNA was extracted from each sample using the RNeasy Plant Mini Kit (Qiagen) following the manufacturer’s instructions and using both the optional QIAshredder columns and on-column DNA digestion. RNA purity and concentration were determined using a NanoDrop One spectrophotometer (Thermo Fisher Scientific) and a Qubit RNA High Sensitivity Assay kit (Thermo Fisher Scientific). cDNA was synthesised with 400 ng RNA and Superscript iv (ThermoFisher) following the manufacturer’s instructions and using the optional RNAse H digestion and oligodT15 primers (Promega). qRT PCR was performed with the LUNA Universal qPCR Master Mix (NEB) following manufacturer’s instructions and 1 µl of cDNA. Data were normalised using the Pfaffl method (Pfaffl 2001) against two reference genes (Hsc_gene_6993 and Hsc_gene_2491). Primers used are available in Table S3. Either One-way ANOVA and Tukey HSD multiple pairwise comparisons or a Kruskal-Wallis test followed by Dunn’s test were performed using R version 4.2.1. The assumptions of normality and variance homogeneity were checked by visual inspection of QQ plots with standardised residuals and residuals versus fitted plots. As an additional criterion the Shapiro-Wilk test and Levene’s test were used. Plots were generated using the ggplot2 v3.4.2 package (Wickham 2016) and figures made in Inkscape v1.1.

#### High Performance Liquid Chromatography (HPLC)

HPLC analysis was performed using a Shimadzu HPLC (Shimadzu Europa GmbH) comprising Nexera X2 binary pump and autosampler with 500 µl sample loop, a Prominence column oven and diode array detector, and fraction collector. The system was controlled using Shimadzu’s Lab Solutions software (version 5.72). Separation was achieved using a YMC-Pack Pro C18 column, 250 mm x 10.0mm ID S-5 µm 12 nm (YMC Europe GmbH Dinslaken). The column was maintained at 40°C and a gradient used for elution at 4.0 ml/min flow rate, with initial composition of 95% mobile phase A (0.1% formic acid) 5% mobile phase B (acetonitrile) changing to 100% B over 16 min, held isocratic at 100% B for a further four min before returning to the initial composition over 1 min and re-equilibrating the column for a further 9 min. 200 µl of sample was injected and fractions were collected every 1 min. Fractions were pooled from a total of eight sample runs and evaporated to dryness. Subsequently, fractions were resuspended in 250 µl ultrapure water.

### Effector network analyses

A transcriptional network of predicted *H. schachtii* effectors was generated as in Molloy et al. (2023) with an arbitrary edge threshold set at a distance correlation coefficient above 0.975. Distance correlation coefficients between the eight Extract upregulated transcription factors (TFs) and predicted effectors were calculated and a network was generated. Of these eight TFs, six were connected to predicted effectors with a correlation coefficient of 0.975 or above. The presence or absence of each predicted effector gene or TF in the Extract upregulated dataset was added to the network as a node attribute. The number of connections with predicted effectors for each TF was added as a node attribute and used to determine the height in the z axis. The network was visualised using Gephi v0.10.1 (Bastian, Heymann, and Jacomy n.d.). Scripts for transcriptional network analyses can be found at: https://github.com/BethMolloy/Effectorome_H_schachtii/tree/main/2ClassNetworkCreation and https://github.com/Jonny-Long-1/The-SUbventral-Gland-master-Regulator-SUGR.

### Characterisation of SUGR1

Domain prediction of SUGR1 (Hsc_gene_14352) was performed using InterPro (Paysan-Lafosse et al. 2023). Protein structure was predicted using AlphaFold (Jumper et al. 2021). The SUGR1 gene model was created using the R package genemodel v1.1.0 (Monroe 2017).

### RNA sequencing and analyses

RNA sequencing and library construction were performed by Novogene. The mRNA library was prepared by poly-A enrichment (poly-T oligo-attached magnetic beads), fragmentation, cDNA synthesis (using random hexamer primers), followed by end-repair, A-tailing, adapter ligation, size selection, amplification, and purification. Illumina sequencing was performed using 150bp paired end reads, generating 5G raw data per sample. RNA sequencing reads are available under ENA accession PRJEB71637. All reads were analysed with FastQC v0.11.9 (Andrews and Others 2010) and 10bp were trimmed using BBduk in BBTools v38.18 (Bushnell 2014). Reads were mapped to the reference *Heterodera schachtii* genome (Siddique et al. 2022) using STAR v2.7.9a (Dobin et al. 2013) and counted using HTseq v0.13.5. (Anders, Pyl, and Huber 2015). Differentially expressed genes were identified in R version 4.2.1 (R Core Team 2022) using the DESeq2 v1.38.3 package (Love, Huber, and Anders 2014) following pairwise comparison of all samples (|log2FC| ≥ 0.5 & padj ≤ 0.001). Hierarchical clustering was performed after scaling using the hclust() function of the stats v4.2.1 package. Volcano plots were plotted using EnhancedVolcano v1.16.0. (Blighe, Rana, and Lewis 2022). GO term enrichment analyses were performed using the gprofiler2 v0.2.1 package (Kolberg et al. 2020). Gene set enrichment was determined by hypergeometric enrichment tests. For the *sugr1* silencing experiment, the same packages were used but with FastQC v0.11.8, BBduk v38.34, STAR v2.7.0e, HTSeq v0.12.4, R v3.5.2, DESeq2 v1.22.2, EnhancedVolcano v1.0.1 and gprofiler2 v0.1.6.

### *In situ* hybridisations

The multiplexed Hybridisation Chain Reaction (HCR) *in situ* was performed as described in (Sperling and Eves-van den Akker 2023). The probes to the designated genes (Hsc_gene_14352; Hsc_gene_2726; Hsc_gene_21727 *eng2*) and *in situ* reagents were designed and purchased from Molecular Instruments, Inc. The images were acquired on a Leica Stellaris 8 FALCON confocal microscope with minor adjustments made to the brightness and contrast. 3D projections were created with the Leica Cyclone 3DR software. The images were prepared using ImageJ (Schindelin et al. 2012). No further image manipulation was performed.

*In situ* hybridisations were performed using ppJ2 of *H. schachtii* following previously published methodology (de Boer et al. 1998). Specific primers were designed to amplify a product for each of the candidate effector genes using a cDNA library produced from ppJ2s (Table S3). The resulting PCR products were then used as a template for generation of sense and antisense DIG-labelled probes using a DIG-nucleotide labelling kit (Roche, Indianapolis, IN, USA). Hybridised probes within the nematode tissues were detected using an anti-DIG antibody conjugated to alkaline phosphatase and its substrate. Nematode segments were observed using a DP73 digital Olympus camera mounted on a Bx51 Olympus microscope.

### Yeast one Hybrid

#### Plasmid construction

Promoters of SUGR-activated genes (up to 2 kb upstream intergenic DNA) were amplified from *H. schachtii* gDNA (Q5 polymerase (NEB) according to manufacturer’s instructions) and cloned into the SacI (NEB) digested pAbAi plasmid (Takarabio). All promoters were additionally analysed in two parts (proximal (b) and distal (a) halves). For this, one promoter half was cut out of the plasmid by mutagenesis PCR (PrimeSTAR Max polymerase (Takarabio) following manufacturer’s instructions). *sugr1, safta,* and *satfb* were amplified from *H. schachtii* cDNA (Q5 polymerase (NEB)) and cloned into the PCR amplified pDEST22 plasmid (Invitrogen). Cloning was performed using the In-Fusion HD cloning master mix (Takarabio) following supplier’s instructions. Bacterial transformation was performed using the heat shock 14 method (30 min ice, 35 sec 42°C, 5 min ice) and plasmids were extracted using the Monarch Plasmid Miniprep kit (NEB) following manufacturer’s instructions.

#### Generation of promoter bait:transcription factor prey yeast strains

Bait yeast strains were generated by transforming the *Saccharomyces cerevisiae* Y1HGold strain (Takarabio) with the promoter (bait) plasmids. Prior to transformation the bait plasmids were linearised in the *URA3* gene by PCR or restriction digest (BbsI/Esp3I/BstBI (NEB)) to allow integration into the genome. Subsequently, the bait strains were transformed with a transcription factor (prey) plasmid to generate bait:prey strains. As a control, all bait strains were also transformed with the pDEST22 empty vector. For yeast transformation, yeast overnight cultures (1.5 ml grown SD-ura or YPDA medium at 28°C) were pelleted and resuspended in 10 µl TE-LiAc solution (10 mM Tris-HCl pH 8.0, 1 mM EDTA, 0.1 M Lithium acetate), 10 µl salmon sperm DNA (Invitrogen), 300 ng plasmid DNA, and 500 µl PEG-TE-LiAc solution (40% PEG 3500, 10 mM Tris-HCl pH 8.0, 1 mM EDTA, 0.1 M Lithium acetate). After shaking at 200 rpm/30°C for 30 min followed by 45 min heat shock at 42°C, yeast were washed with sterile ultrapure water and grown on selective SD medium for three days at 28°C.

#### Yeast-one-hybrid screens

The generated bait:prey yeast strains were grown overnight in liquid SD -ura/trp medium at 28°C and 250 rpm and subsequently diluted in water to OD600 = 0.6. Finally, 2 µl drops of yeast suspension were plated in five 1:5 serial dilutions on 100 mm square plates (Thermo Fisher) with selective SD medium and increasing concentrations (0.07–7 µg/ml) of the antibiotic Aureobasidin A (Takarabio). Yeast were grown for four days at 28°C and images taken on day 2, 3 and 4 using a GBox gel doc system (Syngene). Due to the different background of native transcription factor:bait binding, pictures shown represent the Aureobasidin A concentration and time point at which no or limited growth was observed for the empty vector control yeast strain. Only interactions observed in at least three technical replicates are shown. Pictures were cropped and figures were made using Inkscape but all comparisons shown stem from the same plate. Original pictures are available on request.

### SUG box identification

Proximal 5’ promoter regions of all *H. schachtii* genes were predicted using a series of custom python scripts (https://github.com/sebastianevda/H.schachtii_promoter_regions). In brief, proximal 5’ promoter regions were defined as n bases of intergenic space, where available, upstream of the coding start site. In this study, 800 bp was used. From this database of promoter regions, subsets were extracted and compared. Comparisons included SvG effectors vs DG effectors; and SvG effectors, J2-10hpi expressed genes, or SUGR-regulated effectors vs a random set of 669 genes. Enriched motifs were identified using HOMER (Heinz et al. 2010).

### RNA interference

A silencing mix was prepared using 3 μg/μL dsRNA (either silencing *sugr1* or *gfp*; ordered from Genolution, Table S3); 50 mM octopamine and M9 buffer. *H. schachtii* J2s were soaked in the silencing mix for 48h at 700 rpm on a thermoblock set at 21°C. If needed, silenced J2s were subsequently flash frozen in liquid nitrogen and RNA extraction and sequencing were performed as described in the previous sections.

### Plant penetration assay

Five day old *S. alba* plants (grown on Daishin agar at 21°C) were infected with ∼100 *H. schachtii* J2s (silenced in either *gfp* or *sugr1* as described previously) and kept in the dark for 10h). For staining, roots were treated with 1% bleach for 2 min followed by treatment with boiling acid fuchsin solution for 2 min. Subsequently, roots were covered in acidified glycerol and left to destain. Nematodes counted under a dissecting microscope The results were validated in two independent experiments. Data shown represent three replicates with about 50 plates each. Statistical analysis was performed by a Games-Howell test (Games and Howell 1976; Ruxton and Beauchamp 2008). Adjusted p-value corresponds to the Family Error Wise Rate (Tukey 1953).

### Heterodera glycines SUGR homology

The *H. glycines* SUGR homologue was identified using BLAST (wormbase-parasite (Howe et al. 2017)), and sequence similarity was compared using amino acid alignments in muscle (Edgar 2004). Sense and antisense RNA were synthesised in a single *in vitro* reaction using the MEGAscript® RNAi Kit (Thermo Fisher Scientific, Waltham, MA, USA) according to the manufacturer’s instructions, with an incubation period of 6 hr to enhance RNA yield. The resulting dsRNA product underwent purification (Green and Sambrook 2020), integrity examination through 1.2% agarose gel electrophoresis. Approximately 30,000 freshly hatched *H. glycines* J2s (TN10) per biological replicate were soaked in a mixed buffer containing 3μg/μL dsRNA in 1/4 M9 buffer (43.6 mM Na2HPO4, 22 mM KH2PO4, 2.1 mM NaCl, 4.7 mM NH4Cl), 1 mM spermidine, and 50 mM octopamine at 26 °C on a rotator covered with aluminium foil to maintain a dark environment. After 24 hours of incubation, J2 were washed three times with Nemawash (5 μl of Tween20 in 50 ml of MES buffered water) before being flash frozen in liquid nitrogen if needed.

RNA extraction was performed using the Nucleospin microRNA kit (Macherey-Nagel, Hoerdt, France) following the manufacturer’s instructions. The isolated RNA was then reverse transcribed into first-strand cDNA using LunaScript RT SuperMix (NEB) according to the manufacturer’s instructions. Real-time PCR reactions were conducted using iTaq universal SYBR Green super mix (Bio-Rad) on a CFX96 Real-time PCR Machine (Bio-Rad Laboratories, Inc., Hercules, CA, US) following the manufacturer’s instructions. Thermocycler conditions comprised an initial denaturation cycle at 95 ◦C for 30 s, followed by 40 cycles at 95 ◦C for 5 s and 58 ◦C for 30 s, concluding with amplicon dissociation. The experimental design included three biological replicates and three technical replicates. Expression levels of *Hg sugr* and four subventral gland effectors (Hetgly05367, Hetgly08289, Hetgly20776, and *Hetgly14426*) were normalized to the endogenous *HgGAPDH* (*CA939315.1*) using the Pfaffl method (Pfaffl 2001). Statistical analysis was performed by two-sample t-tests using R version 4.2.1. Plots were generated using the ggplot2 v3.4.2 package and figures made in Inkscape v1.1. RNA sequencing was performed (by Novogene) and analysed as described in previous sections.

*Glycine max* seeds (Williams 82) were surface-sterilized with 70% ethanol for 2 minutes and then with 50% bleach for 10 minutes, followed by three rinses in sterile water. Sterilized seeds were placed on wet filter paper with MES buffer inside a Petri plate and incubated in a growth chamber at 26 °C. 5 days old seedlings were used for the experiments. A 23% Pluronic F-127 (PF-127) (Sigma-Aldrich) gel was prepared as per (Wang, Lower, and Williamson 2009). SCN infection was assessed in a 6-well tissue culture plate. Three milliliters of Pluronic gel were poured into each well, and seedlings were placed in each well at 15-20 °C. After the gel solidified, approximately 100 J2s/50 μL of *H. glycines* were inoculated at the root tip of each seedling using a pipette tip. Nine plates were included in the experiment for each treatment. Three biological replicates were used for each treatment (*gfp* and *Hg sugr*), with each biological replicate consisting of 15 technical replicates (15 individual seedlings). In total, 45 plants for *gfp* and 45 plants for *Hg sugr* were included in the analysis. After 24 hours, plants were harvested from the gel by briefly placing the plates over an ice bath. Due to the slight decrease in temperature, the gel liquefied, allowing the plantlets to be easily extracted without damaging the root system. Roots were stained with acid fuchsin following the method outlined by (Bybd, Kirkpatrick, and Barker 1983), and the number of J2s penetrating the root was counted using a stereomicroscope. Photographs were taken.

## Data and material availability

Raw reads deposited in ENA accession PRJEB71637 Scripts unique to this manuscript are deposited under the following github accessions: https://github.com/sebastianevda/H.schachtii_promoter_regions, https://github.com/Jonny-Long-1/The-SUbventral-Gland-master-Regulator-SUGR Network files are deposited under DRYAD accession DOI: 10.5061/dryad.vmcvdnd0q Plasmids generated are available upon request.

## Author contributions

Data analysis - Clement Pellegrin and Anika Damm

Maintenance of plant-parasitic nematodes - Beatrice Senatori.

Effectostimulins - Anika Damm, Andrea Díaz-Tendero Bravo, Sarah Jane Lynch, Paul Brett, and Clement Pellegrin.

RNAi - Clement Pellegrin

RNAseq - Anika Damm and Clement Pellegrin.

Effector network analyses - Beth Molloy, Dio S. Shin, and Jonathan Long.

*In situ* hybridisation - Paulo Vieira and Alexis Sperling.

Yeast-one-hybrid - Anika Damm, Beth Molloy, Clement Pellegrin

Comparative analyses in *Heterodera glycines* - Joffrey Mejias, Anil Kumar, Rick Masonbrink, Tom Maier, and Thomas Baum.

Wrote the manuscript - Anika Damm, Clement Pellegrin, and Sebastian Eves-van den Akker.

## Acknowledgements

Anika Damm would like to thank Belén Rombolá Caldentey and Sandra Schmöckel (University of Hohenheim) for teaching her the fundamentals of yeast work applied in this study. Anika Damm would like to further acknowledge Victor Hugo Moura de Souza (University of Cambridge) for Bioinformatic support. The authors would like to thank Anna V. Protasio (University of Cambridge) for helpful discussions on the manuscript.

## Declarations

### Ethics approval and consent to participate

Not applicable.

### Consent for publication

Not applicable.

